# Diagnostic utility of telomere length measurement in a hospital setting

**DOI:** 10.1101/225797

**Authors:** Jonathan K. Alder, Vidya Sagar Hanumanthu, Margaret A. Strong, Amy E. DeZern, Susan E. Stanley, Clifford M. Takemoto, Ludmila Danilova, Carolyn D. Applegate, Stephen G. Bolton, David W. Mohr, Robert A. Brodsky, James F. Casella, Carol W. Greider, J. Brooks Jackson, Mary Armanios

## Abstract

Very short telomere length (TL) provokes cellular senescence *in vitro*, but the clinical utility of TL measurement in a hospital-based setting has not been determined. We tested the diagnostic and prognostic value of TL measurement by flow cytometry and fluorescence in situ hybridization (flowFISH) in individuals with mutations in telomerase and telomere maintenance genes, and examined prospectively whether TL altered treatment decisions for patients with bone marrow failure. TL had a definable normal range across populations with discrete lower and upper boundaries. TL above the 50^th^ age-adjusted percentile had a 100% negative predictive value for clinically relevant mutations in telomere maintenance genes, but the lower threshold for diagnosis was age-dependent. The extent of deviation from the age-adjusted median correlated with the age at diagnosis of a telomere syndrome as well as the predominant complication. Mild short telomere defects manifested in adults as pulmonary fibrosis-emphysema, while severely short TL manifested in children as bone marrow failure and immunodeficiency. Among 38 newly diagnosed patients with bone marrow failure, TL shorter than the 1^st^ age-adjusted percentile enriched for patients with germline mutations in inherited bone marrow failure genes, such as *RUNX1*, in addition to telomere maintenance genes. The TL result modified the hematopoietic stem cell donor choice and/or treatment regimen in one-fourth of the cases (9 of 38,24%). TL testing by flowFISH has diagnostic and predictive value in definable clinical settings. In patients with bone marrow failure, it altered treatment decisions for a significant subset.

## Introduction

When primary human fibroblasts are grown in culture, they have a finite replicative potential; it is predictable based on the length of telomere repeat DNA(1). Telomeres define the ends of chromosomes and function to preserve genomic integrity; they are comprised of TTAGGG sequences that are bound by specialized proteins(2). Telomere length shortens during DNA replication, and, at a critical threshold, the shortest telomere(s) activate a DNA damage response that signals cell death or a permanent cell cycle arrest known as cellular senescence(3, 4). The observations in cultured cells, and the fact that telomere length shortens with aging, have led to a hypothesized role for telomere shortening in human aging and age-related disease(1, 5); however, the short threshold that is clinically relevant for disease risk is not known, and whether telomere length measurement can influence treatment decisions in clinical settings has not been examined.

The short telomere syndromes are a group of genetic disorders that are caused by mutations in components of the telomerase enzyme and other telomere maintenance genes(6). They provide a model for understanding the causal role of telomere shortening in human disease. Their manifestations include a heterogeneous spectrum of progressive pathologies that include primary immunodeficiency and/or bone marrow failure(7-9). The most common short telomere syndrome pathology is idiopathic pulmonary fibrosis (IPF) and the related interstitial lung diseases(10, 11). Severe emphysema, alone or combined with fibrosis, may also manifest in short telomere patients who are smokers(12). Mutations in the essential telomerase enzyme genes, *TERT* and *TR,* are the most common cause of IPF, and the frequency of *TERT* mutations in severe, early-onset emphysema rivals the prevalence of alpha-1 antitrypsin deficiency(12).

The prevalence of telomerase mutations in adult-onset lung disease makes the short telomere syndromes the most common of premature aging disorders with an estimated 5,000-10,000 individuals affected in the United States alone(13). Identifying these patients is critical for clinical management as they are susceptible to life-threatening toxicities from DNA damaging agents and other cytotoxic therapies(14-17). In particular, in the hematopoietic stem cell transplant setting for bone marrow failure, reduced intensity regimens improve outcomes(14), yet the vast majority of patients lacks recognizable features at the bedside, and there is often no notable family history at the time of diagnosis(9, 18). There is therefore a need for molecular diagnostic tools for their evaluation at the bedside.

To identify an ideal method for TL measurement in clinical settings, we considered such a method must be reproducible and amenable to standardizing. One method that was developed for epidemiologic studies measures TL by quantitative PCR, but it has been found to have a 20% variability across laboratories(19, 20). This variability lies in part due to its sensitivity to DNA quality and extraction methods, while also being prone to error propagation with PCR amplification(21, 22). Circulating leukocytes also have variable TL depending on their replicative histories(23), thus making the analysis of total leukocyte TL, by quantitative PCR as well as the Southern blotting method, confounded by fluctuations in leukocyte ratios.

Importantly, and in contrast to other genotypes, TL is a continuous variable, and the absence of standardized normal ranges that are clinically relevant presents an impediment to translating research findings to clinical settings. Here, we report a hospital-based experience of TL testing by automated flow cytometry and fluorescence in situ hybridization (flowFISH)(24). This method measures single cell TL using a fluorescently labeled probe that hybridizes to the telomere DNA sequence. We show that peripheral blood TL measurement using flowFISH can inform diagnostic and patient care decisions in definable clinical settings.

## Results

### Telomere length has a discrete and reproducible normal range in the human population

To test the relevance of TL measurement in clinical settings, we sought first to define a normal range based on values obtained from 192 controls across the age spectrum who were recruited at Johns Hopkins Hospital (Table S1). TL shortened with age in lymphocytes and showed a normal distribution at every age, similar to prior reports(11, 23) (Figure 1A, Figure S1A and Table S2). The reproducibility for TL measured by flowFISH was superior to quantitative PCR with an intra- and inter-assay coefficient of variation of 2.2% and 2.5% for lymphocytes, compared to 8.0% and 25.0%, respectively (Figure S1B-D). We compared the inter-lab reproducibility with a Vancouver lab that uses flowFISH and found an outstanding concordance (R^2^ 0.94, slope 1.04, P<0.001, linear regression, Figure 1B and Figure S1B,C). To test the generalizability of this normal range, we compared the nomogram from the Baltimore population with an ethnically different population of 444 controls recruited in Vancouver(23), and found they were nearly identical in range and percentile distributions (Figure 1A). The concordance was tighter for lymphocytes than granulocytes, likely in part because of the known freezing effects on altering granulocyte TL(23) (Figure S1E-F). These data indicated that TL as measured by flowFISH has reproducible and standard lower and upper boundaries across populations.

**Figure 1.**
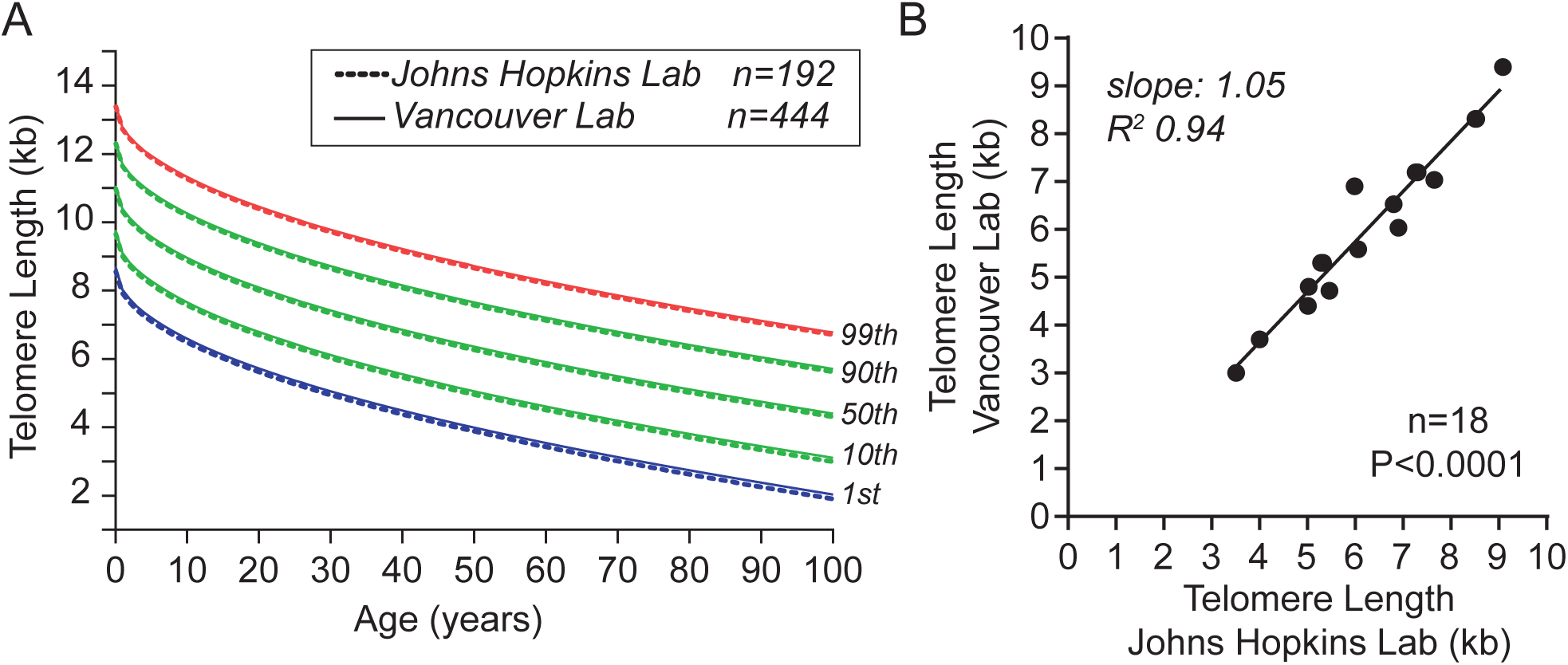
Telomere length by flowFISH shows a reproducible and definable normal range. **A.** Nomogram of lymphocyte lengths from Johns Hopkins controls (n=192) and controls from Vancouver (n=444) with boundaries shown by percentiles as annotated. **B.** Inter-laboratory reproducibility of lymphocyte telomere length measurements from 18 samples, processed independently, shows outstanding concordance by linear regression.

### There is a short telomere threshold associated with disease in humans

We examined whether TL can be used to identify patients with germline defects in telomere maintenance. We recruited 100 individuals from 60 families who carried known pathogenic mutations in telomerase and other telomere maintenance genes (Table S3). Mutations that were deemed pathogenic based on segregation in families and functional molecular evidence fell in seven genes (Table S4): *TERT*, *TR*, *DKC1*, *RTEL1*, *PARN*, *TINF2* and *NAF1* with *TERT* mutations being most common, seen in 49 of the cases. The mean age was 41 (range, 8 months to 77 years) and 54% were male. None of the mutation carriers had TL above the age-adjusted 50^th^ percentile, indicating a 100% negative predictive value for identifying a clinically meaningful mutation (Figure 2A). We examined whether there is a TL threshold, relative to controls, that can identify telomerase and telomere maintenance gene mutation carriers, and found an age-dependent effect. Since the trends we will report hereafter also show for granulocytes, we summarize the analyses for lymphocytes and the respective granulocyte data are in Figure S2. No single threshold identified all mutation carriers in contrast to what has been proposed(25). However, whereas in children under the age of 20 TL deviated significantly from the age-adjusted median control length, adults over the age of 60 usually overlapped with the lowest deciles of controls (mean deltaT -4.4 vs. -1.6 kb, respectively, P<0.0001, Mann-Whitney test, Figure 2A-B). In a similar analysis, 39 of 42 (93%) who were identified before the age of 40 had TL below the first percentile, while only 9 of 58 (16%) diagnosed after 40 years fell in that range (Figure 2B). There was a notable dropout of mutation carriers who had TL below the first percentile after age of 60, possibly reflecting attrition related to more severe telomere-mediated disease (Figure 2A-B). These data indicated that TL testing has an outstanding negative predictive value for excluding individuals with clinically relevant mutations across the age spectrum. In patients younger than the age of 40, the sensitivity is higher.

**Figure 2.**
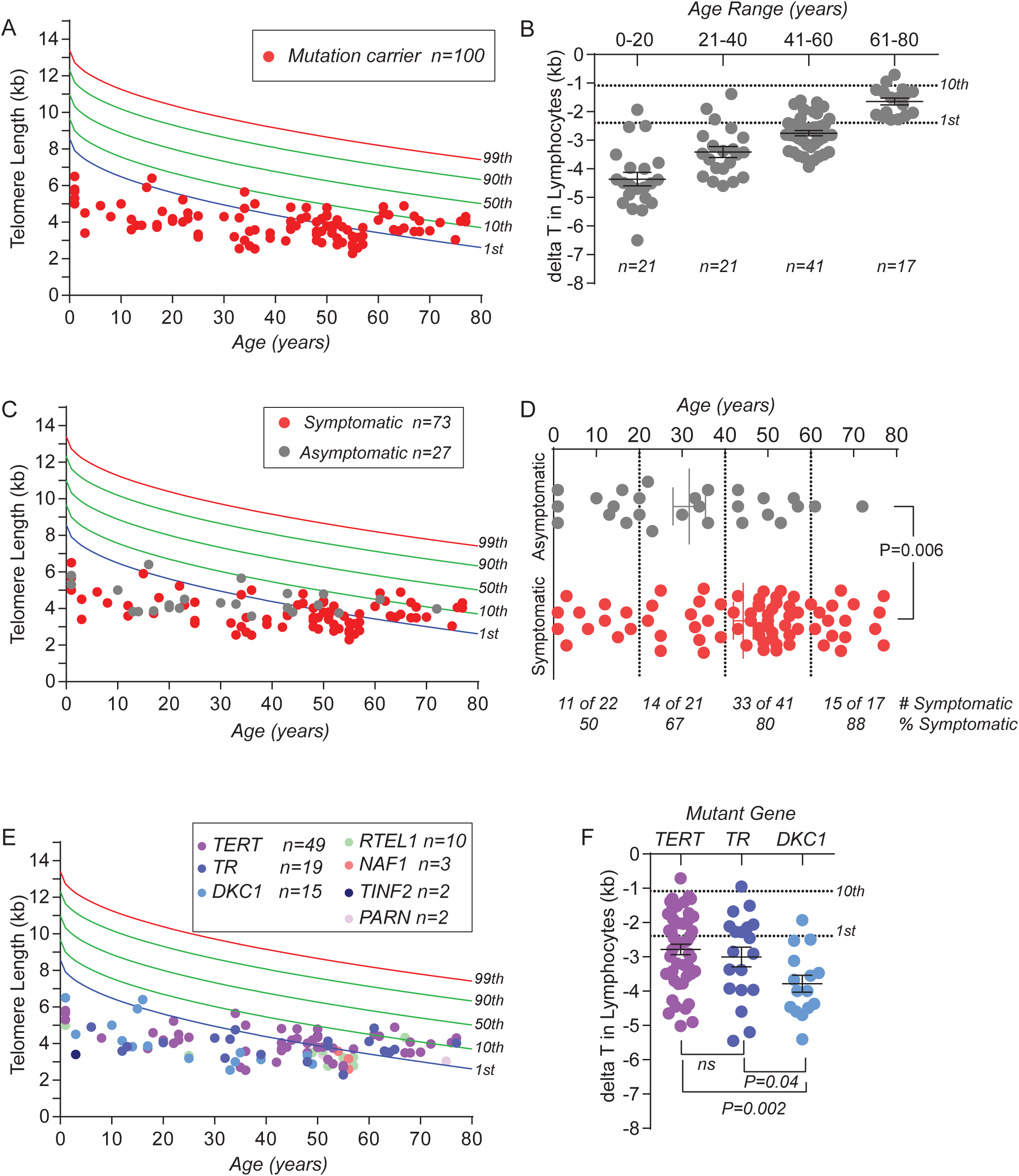
Telomere length has age-dependent diagnostic thresholds. **A** and **B.** Lymphocyte telomere length measurements from 100 telomerase and telomere gene mutation carriers relative to the nomogram with the age-dependent deviation shown by two decade intervals. In B, the 1^st^ and 10^th^ percentile thresholds are annotated to the right. **C.** Data from **A** annotated by mutation carriers who had symptoms, defined as primary immunodeficiency, bone marrow failure, liver disease and/or pulmonary fibrosis-emphysema (left panel), and those who had no symptoms. **D.** Proportion of symptomatic patients increases with age as shown by the average and distribution in the graphic and the proportions affected listed below. **E.** Telomere length in lymphocytes annotated for each patient by mutant gene. **F.** Shorter telomere length in *DKC1* mutation carriers (males) relative to *TERT* and *TR*. Graphs in **B**, **D** and **E** indicate means ± SEM, Mann-Whitney test. Delta T refers to the deviation from the age-adjusted median.

### Telomere length is associated with disease onset and disease type in mutation carriers

We queried the records of these 100 mutation carriers to assess if they had any manifestations of a short telomere syndrome including blood, immune abnormalities, liver and/or pulmonary disease. We found a significant subset was asymptomatic (27 of 100); these patients had been identified because of the diagnosis in a symptomatic relative. The extent of telomere shortening in these asymptomatic individuals was similar to symptomatic patients (mean delta -3.1 vs. -3.0, P=0.7, Mann-Whitney test), suggesting the short telomere defect itself is not pathognomonic of a disease state. However, in this cross-sectional analysis, we noted that the proportion of symptomatic patients increased with age: 15 of 17 (88%) above 60 years showed symptoms compared with 10 of 21 (50%) who were younger than 20 years [(P=0.015, Fisher’s exact test, Figure 2D, Odds Ratio 8.3 (95% confidence interval, 1.5 to 45.4)]. On average, symptomatic individuals were 13 years older than asymptomatic mutation carriers (mean 44.6 vs. 31.7 years, P=0.006, Mann-Whitney test, Figure 2D). There was notably no significant correlation between the mutant gene and TL (Figure 2E), except for *DKC1* mutation carriers (all male) who had shorter TL relative to *TERT* and *TR* mutation carriers (Figure 2E and Figure S2B). Notably, among all 100 mutation carriers, 93 lacked any of the classic mucocutaneous features associated with the short telomere syndrome dyskeratosis congenita highlighting the importance of molecular and the genetic diagnostic tools. The remaining 7 cases were all male and carried mutations in *DKC1*; three had all three mucocutaneous features and four had only one.

We next tested whether in the 73 symptomatic mutation carriers, TL had predictive or prognostic value. We focused on the four common degenerative, fatal complications and noted several significant associations. We found evidence for an age-dependent pattern of telomere-mediated disease, as has been hypothesized(18). Patients who developed bone marrow failure first were on average 33 years younger than pulmonary fibrosis-emphysema patients with normal blood counts (mean 24.5 vs. 57.7 years, P<0.001, Mann-Whitney test). There was also a continuous correlation between the severity of the telomere defect (as measured by the delta telomere length in lymphocytes) and the age at diagnosis (Figure 3B and Figure S2C-D). More severe telomere shortening manifested earlier in life and milder defects provoked disease in adulthood (R^2^ 0.59, slope 13.2, 95% confidence interval 10.6 to 15.8). The correlation between TL severity and disease onset resembled what has been described in mice with short telomeres(26). There was also a correlation between TL and disease complication. Patients with bone marrow failure had significantly shorter age-adjusted TL than patients with lung disease (mean delta -4.0 kb vs. -2.0 kb, P<0.001, Mann-Whitney test, Figure 3C), signifying more severe disease. Patients with liver disease, in contrast, presented at an intermediate age (mean 40.6 years), and had intermediate TL (mean delta -2.8 kb) relative to bone marrow failure and pulmonary fibrosis patients. Although bone marrow failure co-occurred in older adults with pulmonary fibrosis, there was a notable demarcation of lung disease to adulthood, and we found no cases of *de novo* pulmonary fibrosis (i.e. outside of the post-hematopoietic stem cell transplant setting) prior to the age of 35 years (41 of 41 cases, Figure 3A).

**Figure 3.**
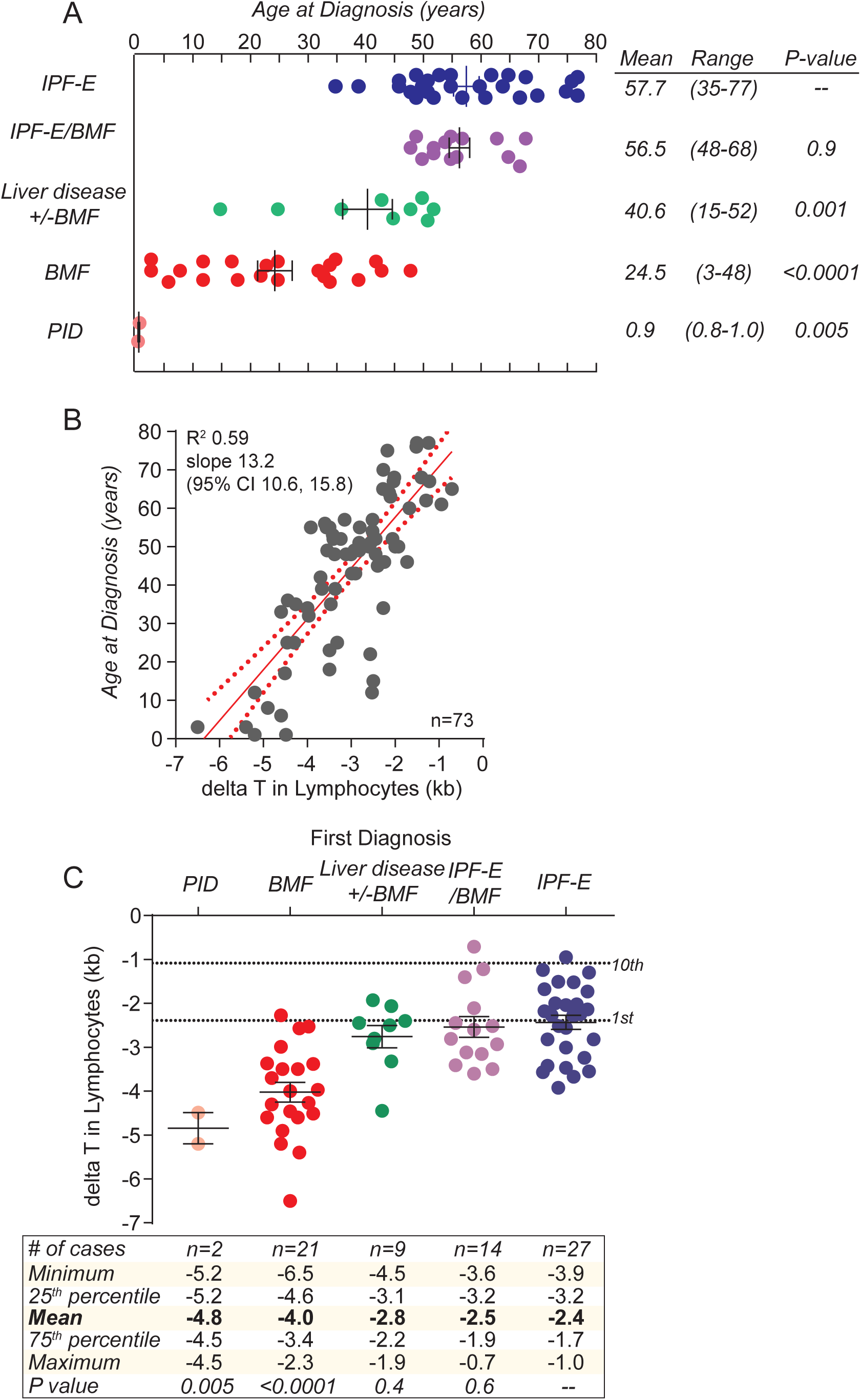
Telomere length correlates with disease onset and type. **A.** Dot plot shows age-dependent manifestations of the four common short telomere syndrome features. IPF-E refers to idiopathic pulmonary fibrosis with or without emphysema. BMF refers to bone marrow failure. PID refers to severe immunodeficiency presenting usually in the setting of enterocolitis in infants. The P-values to the right indicate difference in age relative to IPF-emphysema (Mann-Whitney test). **B.** Linear regression shows a correlation between deviation of lymphocyte telomere length and age at diagnosis of one of four short telomere syndrome features. The linear regression line and 95% confidence intervals are shown for the 73 symptomatic individuals from Figure 2C. **C.** The disease type correlates with the extent of lymphocyte telomere length deviation with the percentile ranges as shown below in the tabulated data. The P-values reflect differences in the mean delta telomere length relative to idiopathic pulmonary fibrosis-emphysema patients. The delta telomere length (delta T) refers to the deviation from the age-adjusted median.

### Telomere length alters clinical management for bone marrow failure patients

We next examined whether TL can distinguish the etiology of aplastic anemia, given the risk of immunosuppression in patients with short telomere-mediated aplastic anemia, and the necessity for reduced intensity regimens in the stem cell transplant setting. We recruited 38 patients under the age of 40 since this was the predominant age at presentation for bone marrow failure of telomere-mediated aplastic anemia (Table S5). The study was observational with results disclosed to the treating clinician in real time. Its endpoints were either a genetic diagnosis or response to immunosuppression. In parallel, we screened for the common inherited bone marrow failure genes (detailed in Methods), and correlated the genetic data with TL. Among 38 recruited patients, 22 reached an informative endpoint with an identifiable genetic cause (n=12 of 38) or response to immunosuppression (n=10) (Figure 4A). All 10 patients who were treated with immunosuppression had lymphocyte TL longer than the age-adjusted 1^st^ percentile and all of them had documented improved in blood counts (partial response n=5, complete response n=5) (Figure 4B). In contrast, all the telomerase or telomere gene mutations had short TL below the first percentile (8 of 8) with mutations identified in *TERT*, *TR*, *DKC1*, *TINF2*, and *RTEL1* (Figure 4A-B). Notably, none of these eight patients had a family history of telomere-mediated disease or any stigmata of a short telomere syndrome at the time of diagnosis highlighting the importance of the molecular diagnosis. In addition to these cases, two other patients, with *LIG4* and *RUNX1* mutations, also had TL shorter than the first percentile, suggesting that this range enriches for other inherited bone marrow failure syndromes. The two patients we identified with *GATA2* mutations had lymphocyte TL in the range of immunoresponders. To test if prior immunosuppression for aplastic anemia may affect TL in the diagnostic setting, we studied aplastic anemia patients who were in remission (median 67 months after treatment initiation, range 14-180, Table S6). Similar to *de novo* aplastic anemia patients who went on to respond to immunosuppression, 11 of 11 of these retrospectively studied patients had TL above the first percentile (Figure 4B). These data suggested that longer lymphocyte TL enriches for immune-mediated forms of aplastic anemia and distinguishes them from short telomere-mediated bone marrow failure. We finally tracked whether TL results altered the management for the patients we recruited. In 9 of the 38 patients we prospectively followed (24%), the abnormally short TL result altered the treatment program by the primary clinician including changing the choice of hematopoietic stem cell donor through an evaluation of TL or genetic causes in relatives, reducing the transplant preparative regimen intensity to avoid undue toxicity in abnormally short telomere patients, or avoiding the use of immunosuppression therapy empiric trials in very short telomere patients.

**Figure 4.**
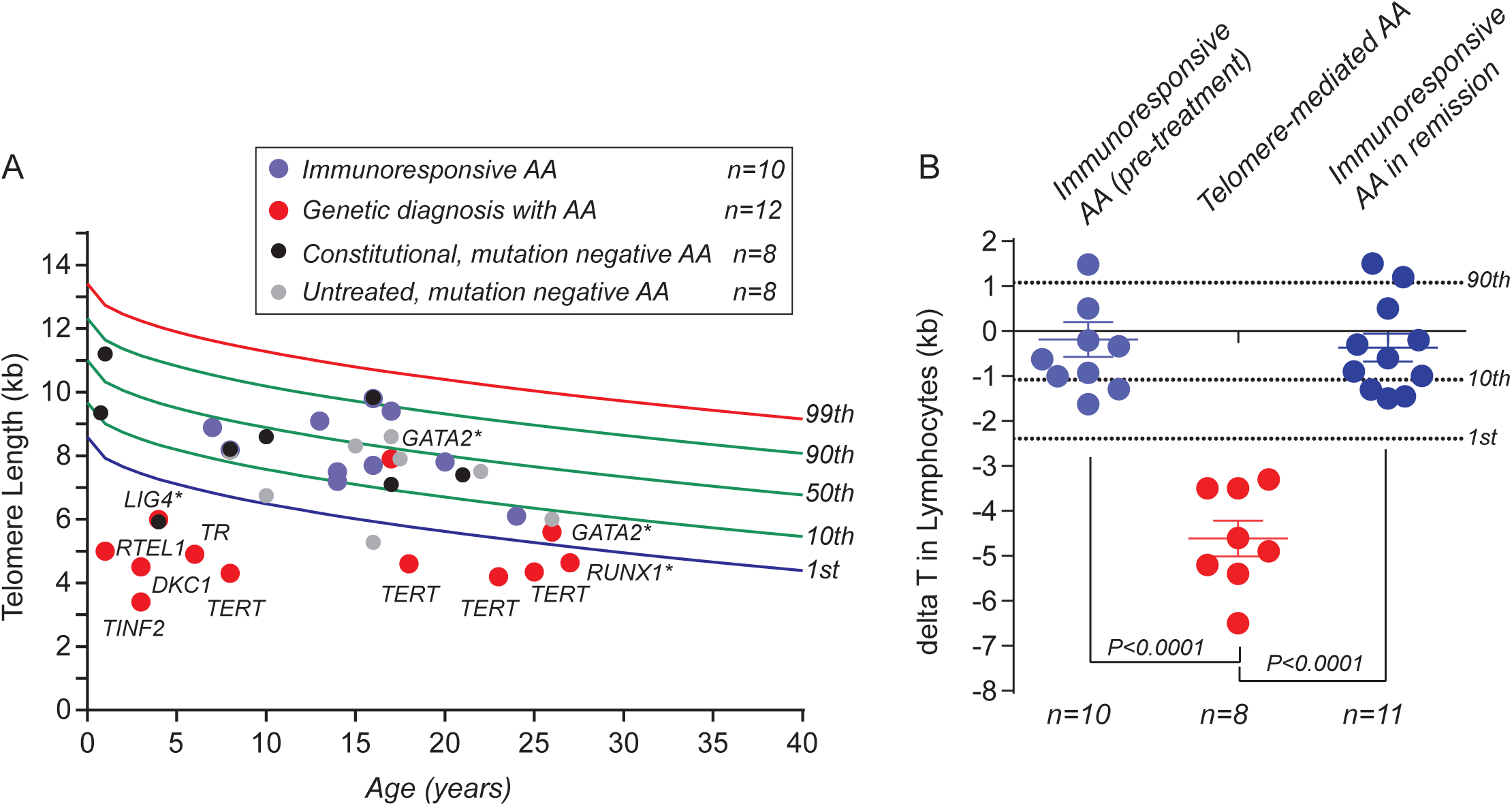
Short telomere length identifies patients with telomere-mediated aplastic anemia in a prospective study. **A.** Lymphocyte telomere length in 38 prospectively recruited patients with aplastic anemia (AA) relative to controls. The red circles denote patients for whom a genetic diagnosis was identified with documentation of a mutation in a telomere maintenance gene (*TERT*n=4, *TR* n=1, *RTEL1* n=1, *DKC1* n=1, *TINF2* n=1) or non-telomere gene (*GATA2* n=2, *RUNX1* n=1, *LIG4* n=1). The latter group is denoted by *. The blue circles denote patients treated with immunosuppression and responded (all treated patients responded). The remaining cases, denoted by black and gray circles, denote cases of constitutional aplastic anemia and untreated cases, respectively. Larger circles indicate the 22 patients who reached an informative endpoint (either a genetic diagnosis made or treatment with immunosuppression). B. delta telomere length (delta T) from the age-adjusted median from three groups: prospectively recruited patients who had a response to immunosuppression at 1 year (n=10, 5 complete response, 5 partial response), and patients with telomerase and telomere gene mutations identified in **A** (*TERT, TR, RTEL1, DKC1, TINF2,* n=8), as well as 11 patients successfully treated with immunosuppression who were in remission for 2 years or more. Means ± SEM are shown, Mann-Whitney test. The 1^st^, 10^th^ and 90^th^ percentile are noted to the right in **B.**

## Discussion

Although telomere shortening has been associated with numerous conditions, its causal role in mediating disease is most strongly linked to genetic disorders caused by mutations in telomerase and other telomere maintenance genes. In our hospital-based study, we found TL by flowFISH had outstanding inter-lab reproducibility, superior to any other TL measurement method heretofore tested. Importantly, we found the flowFISH-derived TL normal range is definable and can be standardized. Given the relatively narrow normal range of TL in the human population, our data indicate that flowFISH is a gold standard for TL testing in patient care settings.

Our study focused on four degenerative disease presentations that are most commonly associated with fatal complications, but patients with mutations in telomerase and telomere maintenance genes may also manifest with enteropathy, infertility and cancer. Short telomere syndromes patients have an increased risk of myelodysplastic syndrome and acute myeloid leukemia as well as squamous cell cancers(27); recognizing these patients has similar implications for management and vigilance for myelosuppression and radiosensitivity with treatment. TL testing provides distinct information from DNA sequencing since it can identify the functional significance of DNA sequence variants. The degree of TL deviation from the median also provides predictive and prognostic information as to the timing of disease onset and the risk of the predominant phenotype likely to develop. The genetic basis of short telomere phenotypes is still uncharacterized in approximately one-third of cases; TL testing may also be a useful diagnostic tool in some of these patients. There are therefore discrete indications for TL testing and the results are critical for diagnostic and prognostic information.

Our data highlight the importance of integrating clinical and genetic findings with TL in diagnostic settings since we did not find a threshold that encompasses all mutation carriers. The first percentile has been suggested to be a cutoff for diagnosis, but we found that would miss a subset of children with this disease and nearly all adult mutation carriers. Instead, older adults with mutations had TL that overlapped with the lowest deciles of the populations though it was always below the 50^th^ percentile and usually below the 10^th^ percentile. While TL in this age group has less diagnostic specificity than in patients younger than 40, the negative predictive value is outstanding. Moreover, the severity of the short telomere defect within this group with predominantly pulmonary fibrosis predicts certain complications (Armanios, unpublished). We also found that the first percentile is not specific and other inherited disorders that cause bone marrow failure may at times be associated with very short TL in leukocytes. The mechanisms are unknown, but it may reflect an acquired shortening related to increased cell turnover, such as with the mutant transcription factor *RUNX1,* or to mutant genes that may possibly affect telomere maintenance, such as with *LIG4.* Regardless, the biology and natural history of these genetic disorders is distinct from short telomere syndromes even though the leukocyte TL may at times be short. Thus, the criteria for the diagnosis of short telomeres syndromes requires integration of TL results with clinical context, and where possible, genetic information. It is important to note that we focused our analyses on lymphocyte and granulocyte TL. We note that although TL measurement in lymphocyte subsets is technically feasible(23), it has failed to show superiority over total lymphocyte TL in distinguishing dyskeratosis congenita from other inherited bone marrow failure syndromes(28).

There has been a recent emerging trend of direct to consumer advertising of TL measurement claiming it may be used to predict biological age and fitness. These methods generally rely on PCR-based quantification which has shown low reproducibility in the literature (19), and in our studies. Our hospital-based experience provides an opportunity to caution clinicians and patients. TL has a significant distribution at every age and small deviations from the median still fall within the normal range and their significance should not be over-interpreted or equated with aging or youthfulness. Rigor in TL measurement is critical as the predicted effect size of changes in TL with some environmental exposures as reported in the literature is often within the error rate of the measurement for the quantitative PCR method(29). Data from recently published large Mendelian randomization studies have identified links between polymorphisms associated with longer TL and the risk of common cancers including lung adenocarcinoma, melanoma, and glioma(30). The upper threshold that increases the risk of these cancers is not known, but these recent findings add significant warning to the over-simplified interpretation of short TL being linked to aging and long telomeres to youth. Overall, the evidence we present suggests that TL testing should be targeted, and that it is most informative in high yield diagnostic settings, such as suspected cases of short telomere syndromes, the interpretation of genetic variation in telomere genes, and in the evaluation of bone marrow failure and related disorders. When a robust and validated method such as flowFISH is utilized in these limited settings, the results have the potential to avert significant morbidity and alter treatment decisions in a way that advances patient care.

## Materials and Methods

### Study review and approval

The clinical studies included here were reviewed and approved by the Johns Hopkins Medicine Institutional Review Board, and all the research subjects gave written, informed consent.

### Healthy controls

To establish a normal range for TL, we recruited 183 volunteers from the Baltimore area across the age spectrum from 2010-2011. Volunteers were interviewed and excluded if they reported having a history of human immunodeficiency virus (HIV), cancer, hemophilia, or pulmonary fibrosis, or if they answered “yes” when asked about a history of a major illness. Race was self-reported as African American, Asian or Caucasian. Peripheral blood was collected in EDTA containing vacutainer tubes (20 cc) (Becton-Dickinson). Cord blood was harvested from 25 discarded placentae. Donor demographics are summarized in Table S1. Mononuclear cells and residual granulocytes were separated by density gradient and frozen at -80°C until analysis. TL measurements were completed by June 2012. Among 208 samples analyzed by flowFISH, 192 (92%) passed preset quality control measures: i.e. less than 10% intra-assay coefficient of variation among triplicate values, and more than 1,000 cell events for each of lymphocyte and granulocyte populations. These values were then used to construct a nomogram as described below. The raw lymphocyte and granulocyte TL values are listed in Table S2.

### Telomerase and telomere maintenance gene mutation carriers

Subjects with mutations in telomerase and telomerase maintenance genes were recruited from 2003-2016 as part of the Johns Hopkins Telomere Syndrome Registry(31-33). Entry criteria for this sub-study to assess the value of TL testing included familial and sporadic cases with a pathogenic mutation in a telomerase or telomere maintenance gene. Rare variants in the known telomere or telomerase genes were deemed pathogenic if they segregated with short telomere phenotypes and/or had functional molecular evidence of pathogenicity (e.g. telomerase activity assay) as previously described(11, 33-36). The mutations are listed in Table S4. Patients who were symptomatic had one of four common short telomere phenotypes: severe combined immunodeficiency, bone marrow failure, liver disease and/or pulmonary fibrosis-emphysema, as described(6). The age at diagnosis and telomere phenotype were extracted from the medical record as previously described(11, 12, 18). Patients with liver disease had hepatopulmonary syndrome or other biopsy-proven cirrhosis. Primary immunodeficiency referred to severe combined immunodeficiency.

### Bone marrow failure study

Newly diagnosed, treatment naïve children and adults with idiopathic bone marrow failure from birth and up to the age of 40 were prospectively recruited from patients who were evaluated at or referred to Johns Hopkins Hospital from 2012-2014. Exclusion criteria included a positive chromosome breakage study, indicating Fanconi anemia, a history of a known inherited bone marrow failure syndrome, personal history of classic mucocutaneous features of dyskeratosis congenita, or a family history of bone marrow failure, pulmonary fibrosis, liver cirrhosis or hematologic malignancy. Patients were classified as having constitutional aplastic anemia if they had a history of developmental delay, intellectual disability or other congenital anomaly. Clinical data were extracted from medical records and study endpoints were assessed at the 2 year timepoint. Table S5 summarizes the clinical characteristics of these 38 patients. Subjects who were retrospectively studied were treated for severe aplastic anemia at Johns Hopkins Hematology Clinics and were recruited from 2011 and 2012 based on a history of durable remission ( >1 year) after immunosuppressive therapy. Their characteristics are summarized in Table S6.

### Telomere length measurement

TL was measured by flowFISH following the detailed protocol in Baerlocher et al.(24), except that PBMCs were extracted by density gradient separation prior to preserving in freezing media. Red cells were lysed at the time of thawing (RBC Lysis Buffer, eBioscience). Cow thymus was obtained from a local butcher and fixed thymocytes were mixed with each sample(24). A peptide nucleic acid (PNA) labeled probe containing the sequence (CCCTAA)_3_ was used for the hybridization (Alexa 488, PNA Bio, California). Each sample was run in triplicate in addition to a no probe to account for background(24). The average lymphocyte and granulocyte TL were then calculated. A reference sample with known TL, obtained from a large leukapheresis product, was used to normalize fluorescence values on each plate. For this sample, the leukocyte TL was measured by Southern blot(1) as well as by flowFISH at a company, Repeat Diagnostics (Vancouver), and the lymphocyte and granulocyte length using this method were found to be similar. Intra- and inter-assay coefficient of variation (CV) was calculated by dividing the standard deviation (σ) by the mean (μ).

### Nomogram generation

Lymphocyte and granulocyte TL from healthy volunteers were read into R/Bioconductor(37). A linear model(38) with the TL as response and square root of age as predictor was fitted based on a strong correlation and supported by evidence of the same relationship in long-lived marine birds(39) (Figure S1A). Using the fitted model, we predicted the TL for individuals 0 to 100 years with the 1 year step. For every predicted value, the 1^st^, 10^th^, 50^th^, 90^th^ 95^th^ and 99^th^ percentile predictions were generated. The same analyses were then performed for the raw control data from Aubert *et al.*(23) (termed Vancouver). Comparisons between the Johns Hopkins and Vancouver labs were limited to similarly prepared samples (i.e. frozen) given the known effect of freezing on TL(23).

### DNA sequencing and analysis

DNA was extracted from peripheral blood (PureGene Blood Core Kit, Qiagen). Mutations were detected by either PCR amplification and Sanger sequencing as described(11, 40), whole exome sequencing(35, 36), whole genome sequencing(33), or TruSeq Custom Amplicon sequencing followed by confirmation with Sanger sequencing(12). For targeted panel sequencing, we designed a TruSeq Custom Amplicon probe set (Illumina) that included the coding and flanking intronic sequences of telomere genes (*TERT*, *TR*, *DKC1*, *RTEL1*, *NAF1*, *TINF2*, *CTC1*, *PARN*, *NOP10*, *NHP2*, *TCAB1*) and *GATA2* (including the intron 4 enhancer). For TruSeq sequencing, libraries were generated from 500 ng of DNA according to manufacturer recommendations and analyzed on a MiSeq sequencer (Illumina)(12). The mean coverage of target sequence was 238x and 88% of the target sequences were covered at or greater than 8x.

### Statistics

We used GraphPad Prism (La Jolla, CA) to graph the data and analyze the statistics, except for the nomogram generation and comparison of distributions which were performed as detailed above. All P-values are two-sided.

## Acknowledgements

We are grateful to patients and study volunteers and to their referring clinicians. We acknowledge support from the Genetic Resources Core Facility staff. We are grateful for helpful discussions with Michael Ochs and Alan Scott and technical support from Laura Kasch-Semenza and Armanios lab members. This work was supported by NIH grants K99-R00 HL113105 (JKA), K23 HL123601 (AED), R37 AG009383 (CWG), and RO1 CA160433 and RO1 HL119476 (MA), the Maryland Stem Cell Research and the Commonwealth Foundations, Johns Hopkins’ inHealth initiative, the Flight Attendants Medical Research Institute and the Gary Williams Foundation (MA). SES received support from NIH T32 GM007309.

## Author Contributions

JKA, CWG, JBJ, MA conceived the idea. JKA and MA designed the research. JKA, VSH, MAS, SES, LD, DWM, JBJ, MA performed research. AED, CMT, LD, CDA, SGB, RAB, JEC, CWG contributed new reagents or analytic tools. JKA, SES, CWG, MA analyzed data. MA wrote the paper. All the authors reviewed and gave comments on the manuscript.

